# Genetic diversity of potato leafroll virus is shaped by variant displacements and selective pressures imposed by aphid and tuber transmission routes

**DOI:** 10.1101/2025.08.18.670828

**Authors:** Graham H. Cowan, Catherine Thomson, Emma Back, Lesley Torrance, Christophe Lacomme, Eugene V. Ryabov

## Abstract

Potato leafroll virus (PLRV) is a major pathogen affecting potatoes worldwide. Since 2018, PLRV incidence increased in Scottish potato crops. Deep sequencing of PLRV in Scottish potato plants revealed the prevalence of a novel PLRV type which became predominant in 2023, displacing the phylogenetically distinct variants that have been present in the region since at least 1989. Analysis of the infection dynamics of the cDNA clone-derived PLRV isolates in potato plants indicated that the novel PLRV may accumulate to higher levels compared to the historic one. Analysis of the genetic diversity of PLRV in early and late field generations (FGs) of seed potatoes showed a significantly reduced genetic diversity of the PLRV structural genes in the early FGs, compared to the late FGs, while divergency of the non-structural genes remained similar across all FGs. Considering that late FGs are more likely to be infected with PLRV via tuber transmission, and early FGs via aphid transmission, these findings suggest that aphid transmission imposes a genetic bottleneck on the structural genes of PLRV, but not on its non-structural genes.

## 1. Introduction

Potato leafroll virus (PLRV) is a positive strand RNA virus which belongs to the species *Polerovirus PLRV* within the family *Solemoviridae* [1-4]. PLRV is a major pathogen of potato (*Solanum tuberosum L*.) which is present in all potato growing regions worldwide and causes significant losses of yield and quality of potato tubers [5-7]. PLRV has a 5.8 kb positive-strand genomic RNA which contains a VPg protein covalently linked to the 5’ terminus. The PLRV genome contains eight open reading frames (ORFs), some of which overlap. The viral RNA replicase is encoded by ORF1 and ORF2, the structural genes are encoded by the ORF3 and ORF5, suppressor of RNA silencing and phloem movement protein – by ORF0 and ORF4, respectively, Expression of PLRV ORFs involves translational frameshift, readthrough, leaky scanning and ribosome scanning strategies [5]. The 5’ proximal ORFs 0, 1, 2 and 8 are expressed from the genomic RNA, while expression of the 3’ proximal ORFs 3a, 3, 4, 6, and 7 occurs from two subgenomic (sg) RNAs sharing the same 3’ as the genomic RNA [4,5]. The larger sgRNA is required for translation of the ORF3, ORF3a, ORF4, and ORF5. The C-terminal 125 amino acid (AA) portion of the ORF5 encoded protein could be expressed as the ORF7 protein from the shorter sgRNA [5].

PLRV forms non-enveloped icosahedral virus particles, approximately 24 nm in diameter, composed of 180 structural protein subunits, the majority of which are coat protein (CP) subunits encoded by the ORF3. During PLRV infection, the readthrough protein (RTP) which contains the ORF3 encoded CP fused to the ORF5-encoded readthrough domain (RTD), is produced as a result of translational readthrough of the amber codon of the ORF3 [4]. The RTP subunits are incorporated into PLRV virus particles as a minor structural protein, with the N-terminal CP part interacting with other CP subunits to form the virion shell, with the RTD protruding from the surface of the virion [8]. The N-terminal part of RTD (^N^RTD) encoded by the ORF5 plays a key role in aphid transmission [9]. The C-terminal part of the RTD (^C^RTD) is proteolytically processed at multiple sites and several truncated RTD variants are included in the PLRV particle [10].

Like other members of the genus *Polerovirus*, PLRV is a phloem-limited virus, which replicates and moves only in the cells of the phloem. The systemic movement of PLRV within plants requires structural proteins encoded by ORF3 and ORF5, as well as movement protein encoded by the ORF4 [4,5]. The systemic movement through the phloem of potato plants also allows PLRV infection to reach developing tubers enabling vertical transmission of the virus to the next vegetative generation. The virus is transmitted between plants by aphids, principally by *Myzus persicae*, in a non-replicative persistent manner [4,5]. Virion assembly is essential for both phloem movement and for the aphid transmission of PLRV. Both the CP and minor RTD are required for the systemic movement of PLRV through the phloem and aphid transmission, which involves transition of the virus particles through midgut and salivary gland epithelial cells and circulation in the hemolymph [9,11]. Interactions with phloem and aphid factors may exert different selective pressures on the structural proteins of PLRV, genetic composition of PLRV populations may be affected by aphid and tuber transmissions.

Although PLRV incidence in Scottish potatoes remain low, PLRV incidence increased since 2018 [12], which could be explained by the higher abundance of aphid vectors of the virus due to the recent ban on the pesticides used for aphid control [13], or other environmental or cultural factors. It is also known that virus epidemics could be a result of the introduction and spread of a novel virus variant(s) [14]. To test this possibility, we analyzed the genetic diversity of PLRV in potato crops using high-throughput sequencing. Our analysis of symptomatic seed potato samples collected across Scotland in 2023 showed the prevalence of a single novel phylogroup of PLRV which apparently had displaced isolates of different phylogroup present in Scotland since at least 1989 [2]. We also investigated the hypothesis that genetic changes in the PLRV populations might have been a result of the virus adaptation to the aphid and tuber transmissions routes by deep-sequencing of PLRV populations from different field generations (FGs) of seed potatoes, which might have acquired PLRV predominantly either from the aphid vectors or vertically from infected tubers. Further, we engineered an infectious cDNA clone of the contemporary Scottish PLRV variant and compared infection dynamics of the clone-derived Scottish PLRV isolates from both phylogroups in potato plants. While this approach does not represent a natural mode of infection, preliminary data suggest a higher accumulation of the PLRV isolate of the currently prevalent phylogroup. Further analyses are needed to conclude whether the prevalent PLRV type has a selective advantage over historic PLRV isolates in natural infection context.

## 2. Materials and Methods

### 2.1. PLRV samples

To analyze the genetic diversity of PLRV in seed potato crops, leaf samples from individual potato plants (n = 889) of 2^nd^ to 6^th^ field generations (FG2 to FG6) showing putative symptoms of PLRV infection were collected across Scottish seed potato growing areas in the summer 2023. Initial screening for the virus was carried in support of the Scottish Seed Potato classification scheme by Science and Advice for Scottish Agriculture (SASA). The presence of PLRV infection was identified by ELISA tests [15]. A representative set of PLRV-infected samples from different FGs was selected for deep sequencing (Table 1). In addition, ware potato plants displaying leafroll symptoms were collected in the summer of 2024 to analyze PLRV circulating in Scotland in the following growing season. This material was used as source in the subsequent production of an infectious cDNA clone (Table 1). To further characterize isolates of the virus in potato crops from previous years, we sequenced PLRV from the potato plants which originated from PLRV-infected seed potato plants collected from seed potato fields in Scotland in 2010, which were then tuber-propagated in insect-proof glasshouse every year until sampling (Table 1).

**Table 1.** PLRV samples used in this study.

| Sample ID | Field sampling year | Region of Scotland | Location | Field Generation | GenBank Accession Number<sup>1</sup> |
| --- | --- | --- | --- | --- | --- |
| 2023_18_W | 2023 | East Central | Forfar | FG6 | PP584529 |
| 2023_19002 | 2023 | Northeast | Inverurie | FG3 | PP584530 |
| 2023_19021 | 2023 | Northeast | Inverurie | FG2 | PP584531 |
| 2023_19056 | 2023 | East Central | Perth | FG6 | PP584532 |
| 2023_19078 | 2023 | Northeast | Inverurie | FG6 | PP584533 |
| 2023_19018 | 2023 | Northeast | Inverurie | FG2 | PP584534 |
| 2023_19027 | 2023 | Southeast | Galashiels | FG5 | PP584535 |
| 2023_19083 | 2023 | Northeast | Inverurie | FG6 | PP584536 |
| 2023_19051 | 2023 | East Central | Perth | FG3 | PP584537 |
| 2023_19074 | 2023 | Northeast | Inverurie | FG2 | PP584538 |
| 2023_19093 | 2023 | East Central | Perth | FG6 | PP584539 |
| 2023_19033 | 2023 | East Central | Perth | FG6 | PP584540 |
| 2023_18997 | 2023 | Northeast | Inverurie | FG5 | PP584541 |
| 2023_18983 | 2023 | Northeast | Inverurie | FG3 | PP584542 |
| 2023_18953 | 2023 | East Central | Perth | FG4 | PP584543 |
| 2023_18957 | 2023 | East Central | Perth | FG5 | PP584544 |
| 2023_18956 | 2023 | East Central | Perth | FG5 | PP584545 |
| 2023_18992 | 2023 | Northeast | Inverurie | FG6 | PP584546 |
| 2023_18980 | 2023 | East Central | Perth | FG6 | PP584547 |
| 2023_MarAG1 | 2023 | East Central | Fife | FG5 | PP584548 |
| 2023_MarAG2 | 2023 | Northeast | Aberdeen | FG3 | PP584549 |
| 2010_1_Es | 2010<sup>2</sup> | East Central | Gogarbank | Ware | PQ047449 |
| 2010_2_K | 2010<sup>2</sup> | East Central | Gogarbank | Ware | PQ047450 |
| 2010_3_BX | 2012<sup>2</sup> | East Central | Gogarbank | Ware | PQ047451 |
| 20104_KF | 2010<sup>2</sup> | East Central | Gogarbank | Ware | PQ047452 |
| 20105_EX | 2010<sup>2</sup> | East Central | Gogarbank | Ware | PQ047453 |
| 2010_EX | 2010<sup>2</sup> | East Central | Gogarbank | Ware | PQ047454 |
| 2024_cDNA_clone | 2024 | East Central | Forfar | Ware | PQ868077* |
| 2024_Forfar_1 | 2024 | East Central | Forfar | Ware | PV647838** |
| 2024_Forfar_2 | 2024 | East Central | Forfar | Ware | PV647837** |

<sup>1</sup> GeneBank accession numbers of de novo assembled contigs, cloned cDNA (marked with \*) or RT-PCR amplicon (marked with \*\*);
<sup>2</sup> Propagated in insect-proof greenhouse RNA was extracted in 2019 (for 2010\_1\_Es), 2018 (for 2010\_2\_K), and in 2023 (for the rest of the samples).

### 2.2. High througput sequencing of PLRV

Total RNA was extracted from potato leaf material using NucleoSpin RNA plant kit (Macherey-Nagel). Reverse transcription was carried out using LunaScript (New England Biolabs) with PLRV-specific reverse primers PLRV-4158REV (5’- CTGAAGGATCCTGCGGTATC-3’) and PLRV-REV (5’-ACTACACAACCCTGTAAGAG-GATCC-3’) to produce cDNAs for the 5’ and 3’ section of the viral genome respectively. Reverse transcription (RT) was carried using LunaScript (New England Biolabs) with specific reverse primers, PLRV-4158-REV (5’- CTGAAGGATCCTGCGG-TATC-3’) and PLRV-REV (5’-ACTACACAACCCTGTAAGAGGATCC-3’) to produce cDNAs for the 5’ and 3’ sections of the viral genome respectively. Two overlapping RT-polymerase chain reaction (RT-PCR) fragments, 4.2 kb and 2.7 kb, covering the entire 5.8kb PLRV genome were amplified with proof-reading Phusion polymerase (New England Biolabs). The 5’ section was amplified with the primers PLRV-FOR (5’-ACAAAAGAATACCAGGAGGAATTGC-3’) and PLRV-4158-REV, the 3’ genome section was amplified with the primers PLRV-3115FOR (5’-GGAG-TATAAAACACTAGGTTTCAAAG-3’) and PLRV-REV. The RT-PCR fragments were isolated from agarose gel with Gel Extraction Kit (QIAgen

High-throughput sequencing was done at the James Hutton Institute Genomic facility. The 5’ and 3’ RT-PCR fragments derived from each sample were pooled equimolarly, quality control of the amplicon pools was analyzed using Bioanalyzer 100 using recommended procedure, the final pools were diluted to 6 pM containing 20% PhiX control DNA and loaded on Illumina MiSeq with paired end 150 bp using standard parameters. Post run fastq files were generated, samples de-multiplexed, and adapters removed onboard for each sample. Libraries of approximately 25 thousand paired-end 150 nt reads were produced for each sample. Cleaned data from each library were then used for *de novo* assembly of the viral contig sequences using rnaSPAdes v3.15.4 (standard k-mers: 21 nt), [16]. The de novo assembled contigs corresponding to the full-length PLRV genomic RNAs were deposited to NCBI GenBank, accession numbers PP584529 to PP584549, and PQ047449 to PQ047454 (Table 1).

To characterize PLRV in the symptomatic samples collected in the year 2024, two overlapping PLRV RT-PCR amplicons were produced as above and then sequenced by using both the Sanger method and the Oxford Nanopore technology with Eurofins Genomics Germany GmbH. The sequences were submitted to NCBI GenBank, accessions numbers PV647837, and PV647838 (Table 1).

### 2.3. Bioinformatic and statistical analyses

The full-length contigs open reading frames (ORF) of PLRV sequences were analyzed and compared against sequences deposited to GenBank by the BLAST search tool [17]. The full-length sequences of PLRV genomic RNA produced in this study and all full-length genomic sequences of PLRV available in NCBI GenBank (n= 87) were aligned with ClustalW [18] (Galaxy Version 2.1+galaxy1) tool and the alignment was used to produce maximum likelihood (ML) tree using RAxML tool (Galaxy Version 8.2.12+galaxy1) using Galaxy server. Bootstrap values were obtained from 100 replicates with PHYLIP package [19]. The trees were visualized by using Figtree tool (http://tree.bio.ed.ac.uk/software/figtree/).

Identification of codons under both stabilizing and diversifying selection was carried out by estimating the relative rates of synonymous (dS) and non-synonymous substitutions (dN) in the PLRV infecting seed potato in the year 2023, by using web based Datamonkey tools (http://www.datamonkey.org) [20].

Pairwise distances between sections of de novo assembled PLRV contigs were calculated by Clustal W [18]. Statistical analyses were conducted using R Software version 4.2. and figures were produced using the packages ggplot2 [21].

### 2.4. Design and evaluation of PLRV cDNA clones

Construction of the full-length infectious cDNA clone of a typical contemporary Scottish PLRV isolate was carried out essentially as previously [22]. Total RNA was extracted from potato plants showing PLRV symptoms sourced in July 2024 in Forfar, Scotland, and was used to produce cDNA with Lunascript RT Supermix Kit (New England Biolabs) and reverse primer PLRV-R2 (5’-<u>TGGA- GATGCCATGCCGACCCA</u>CTACACAACCCTGTAAGAGGATCC-3’) which was complementary the 3’ terminus of PLRV genomic RNA. The cDNA was used as a template to produce PCR fragment using the proof-reading Phusion High Fidelity DNA polymerase (New England Biolabs), and primers PLRV-R2 and PLRV-F2 (5’- <u>GGAAGTTCATTTCATTTGGAGAGGA</u>CAAAAGAATACCAGGAGGAATTGC-3’). The PLRV amplicon was cloned into a vector pDIVA (GenBank Accession Number KX665539), based on the mini binary vector pCB [23], supplemented with a cauliflower mosaic virus (CaMV) 35 S promoter followed by a hepatitis delta virus (HDV) ribozyme and the polyadenylation signal of CaMV. The vector was amplified by inverse PCR using primers pDIVA-F (5’-CCTCTCCAAATGAAATGAACTTCC-3’) and pDIVA-R (5’-GGGTCGGCATGGCATCTCCA-3’). Ligation of the vector and PLRV cDNA amplicons was performed using the NEBuilder HiFi DNA kit (New England Biolabs), standard molecular cloning techniques were used to produce the full-length PLRV clone pDIVA-PLRV-Scot-2024. The plasmid clone was sequenced by both Sanger method and the Oxford Nanopore technology with Eurofins Genomics Germany GmbH, the sequence was submitted to NCBI GenBank, accessions number PQ868077 (Table 1). pDIVA-PLRV-Scot-2024 was transformed into Agrobacterium tumefaciens strain LB4404. Inoculation of potato plants (cv Desiree) with cDNA clone-derived PLRV was carried out using agrobacterium culture method involving incubating the freshly cut stem section with the agrobacterium cultures with PLRV cDNA clones [22]. The plants were rooted in pots containing compost and were grown in a glasshouse with 16 h light (18 °C) and 8 h dark (15 °C).

The levels of PLRV RNA in potato plants were quantified by RT-qPCR (reverse transcription quantitative PCR). Total RNA was extracted as above, the first strand cDNA synthesis was carried using LunaScript (New England Biolabs) with random primers, qPCR was carried out with a SYBR green kit with the primers qPLRV-REV (5’-GCACAGAGAATACTGCAGCCAG-3’ and qPLRV_FOR (5’- TTCAGAAATCCGACCCTCGC-3’) for PLRV RNA target, and the primers qActin_rev (5’-TGATTTGAGTCATCTTCTCACG-3’) and qActin-FOR (5’-GAGCAATTGGGATG ACATGG-3’) for the potato beta-actin mRNA target. The cycle threshold (Ct) values for PLRV were subtracted from the Ct values for the beta-actin to obtain normalized DCt values were used to compare loads of PLRV genomic RNA in potato tissue samples.

Phylogroup identity of PLRV in RNA samples from tuber-propagated potato plants was determined by RT-PCR using primers pairs to the region coding for the C-terminal part of RTD specific either to PLRV type N (5’-CGAAAGCCGATCTTTTA-GAA-3’, and 5’-CAAGCGCTCTTCATAAGTCTGT-3’) or the PLRV type O (5’-CGAAA-GCCGATCTACTGGAG-3’ and 5’-TAAACGCTCTTCAACAGTTACC-3’), the amplified RT-PCR fragments were sequenced by Sanger.

For the expression of the structural proteins of PLRV of the historic and novel Scottish types tagged with the green florescent protein (GFP) tagged, the ORF3-ORF5 coding sequences were amplified by PCR from the infectious cDNA plasmid constructs (GenBank accession numbers OK245432 and PQ868077, respectively) with the primers AttB1-PLRV-P3P5-for (5’-GGGGACAAGTTTGTACAAAAAA-GCAGGCTCGATGAGTACGGTCGTGGTTAAAGGAAATGTCAATGGTG-3’) and AttB2-stp-PLRV-REV (5’-GGGGACCACTTTGTACAAGAAAGCTGGGTttagttaAC-TACACAACCCTGTAAGAGGATCC-3’) and cloned, using Gateway cloning approach, into the binary plant expression vector pGWB406 to produce in frame GFP fusion for expression under the control of the CaMV 35S promoter.

## 3. Results

### 3.1. Identification of a prevalent PLRV phylogroup which became predominant in Scottish potatoes

The genetic diversity of PLRV across all Scottish potato growing areas in 2023 (Table 1) was determined. For the analysis, we sampled individual potato plants of different field generations (FGs) of seed potatoes. To obtain additional sequenced data on the PLRV variants circulating in Scotland previously, we analyzed PLRV genomes in potato plants originating from the tubers collected in the field in 2010 and propagated by tuber transmission in an aphid-free glasshouse until sampling in 2018, 2019 or 2023. Furthermore, we analyzed several samples of ware potatoes in the year 2024. We produced RT-PCR amplicons covering entire PLRV genome, which were sequenced using high-throughput sequencing approach. The sequencing libraries were used to produce the full-length de novo assembled contigs.

The Scottish seed potato production starts from *in vitro* propagated virus-free micro-plants to produce mini-tubers that can be grown in the field to produce progeny tubers (field generation one; FG1). The field-grown progeny tubers are harvested and planted for up to 6 consecutive years to multiply seed tubers output. PLRV could be introduced to potato plants by aphids and then transmitted via tubers. Therefore, the assumption was made that analysis of the samples collected in a single growing season from early and late FGs could give a snapshot of PLRV populations most likely transmitted by aphids in crops of earlier FGs and up to 5 previous years of tuber propagation for crops of later FGs.

Phylogenetic analysis of the nucleotide (NT) sequences of PLRV identified in our study and all full-length PLRV sequences available in GenBank showed that all Scottish seed potato samples collected in the years 2023 and 2024 (Table 1) were infected with the PLRV variants phylogenetically distinct from those collected in Scotland from 1989 to 2021, including PLRV variants originated from the 2010 samples which were sequenced in this study (Fig. 1). The genomes of all Scottish PLRV sequences identified in the 2023 and 2024 form a separate monophyletic clade with 89% bootstrap support, suggesting a monophyletic origin and perhaps recent diversification (Fig. 1, red branches). These sequences were similar to the PLRV isolate identified in Germany in 2012 (GenBank accession JQ346189) [24]. The Scottish PLRV isolated in 2023 and 2024, the German 2012 variant, and three Columbian isolates identified in 2015-2021 clustered together to form a clade with 100% bootstrap support, which we hereafter refer to as PLRV type N (Fig. 1).

**Figure 1.**
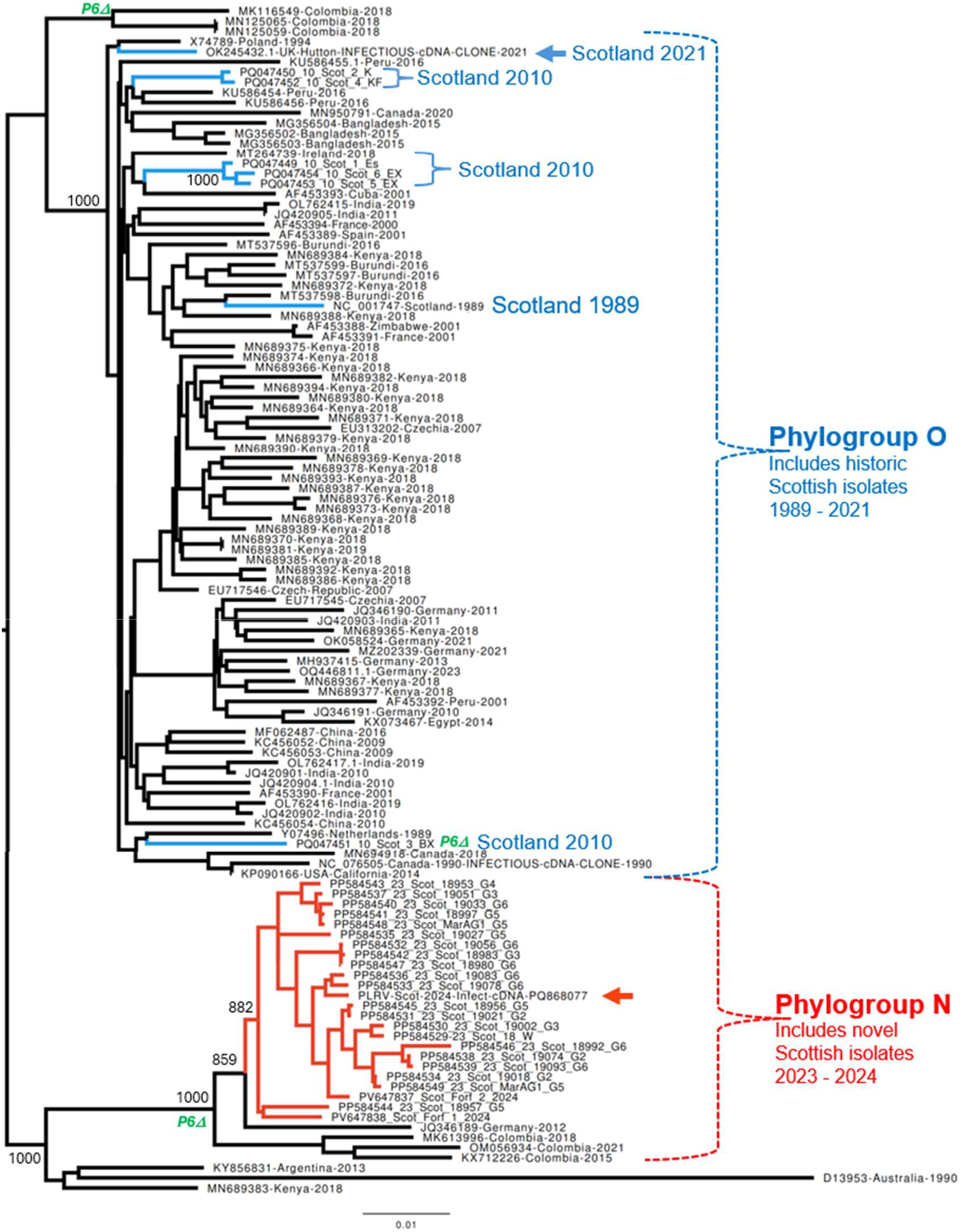
Sequence analysis of PLRV variants identified in Scotland and worldwide. Maximum likelihood (ML) phylogenetic tree for full-length nucleotide sequences of PLRV produced in this study (n = 30) and for all full-length PLRV genomes available in GenBank sampled globally (n = 87). Blue branches indicate Scottish PLRV samples collected from 1989 to 2021, red branches indicate all PLRV samples collected in Scotland during 2023 and 2024. Bootstrap values obtained after 1,000 re-samplings are shown for selected nodes. PLRV variants with truncated P6 are indicated with green “P6D” at the node label or at the base of branch for the groups sharing this feature. The node labels include Gen-Bank accession numbers, geographic location, and the year of sampling.

All the full-length PLRV genomes of Scottish isolates sequenced from 1989 to 2021, including contigs from PLRV infected plants originated from the 2010 field collected plants which were tuber-propagated in aphid proof glasshouse until 2018 - 2023 (Fig. 1, blue branches), belonged to a phylogenetic clade with 100% bootstrap support, which also included PLRV sequences from all potato growing areas worldwide. The Scottish PLRV isolates from this clade, which we refer to as the PLRV type O (Fig. 1), also included a 2021 isolate for which an infectious cDNA clone was constructed [22]. We further tested the presence of both PLRV types in the RNA extracts isolated from Scottish potato samples field-collected in 2019, 2021 and 2022 by strain-specific RT-PCR amplification and sequencing of a 360 nt section of the ORF5. This analysis showed that both the PLRV types O and N were present in Scotland in 2019 (Supplementary Fig. S1, Text S1) suggesting gradual variant replacement.

Figure 2a exemplifies the genetic differences between the PLRV types O and N by comparing the full-length infectious cDNA clones of the PLRV type O (isolated in Scotland in 2021 [22]) and PLRV type N (isolated in Scotland in 2024). The sequences show overall 4% NT divergency, resulting in 82 divergent AAs in the putative proteins encoded by the ORFs 0 to 8. The highest sequence variation was observed in the 3’-parts of the ORF5 coding for the C-terminal, unstructured part of the readthrough protein (RTP), which is also expressed as the P7 protein. Several AA changes was found in the ORF0 protein (P0) which functions as a viral suppressor of RNA silencing (VSR) [5], and in the P1 protein, a part of RNA replicase (Fig. 2a). Recently, it was experimentally demonstrated that the P0, P1 and P7 proteins of PLRV alter interaction between aphids *and Nicothiana benthamiana* plants by modulating production of phytohormones affecting aphid fecundity and attractiveness to PLRV infected plants [25]. A noticeable feature of all PLRV isolates of the Scottish phylogroup N, which was also found in the German 2012 isolate and three South American isolates, was the stop codon in the ORF6 short-ening the P6 protein from 62 to 39 AAs. Such truncated ORF6 was found only in 5 out of 89 full-length PLRV sequences outside of the PLRV type N (Fig. 1).

**Figure 2.**
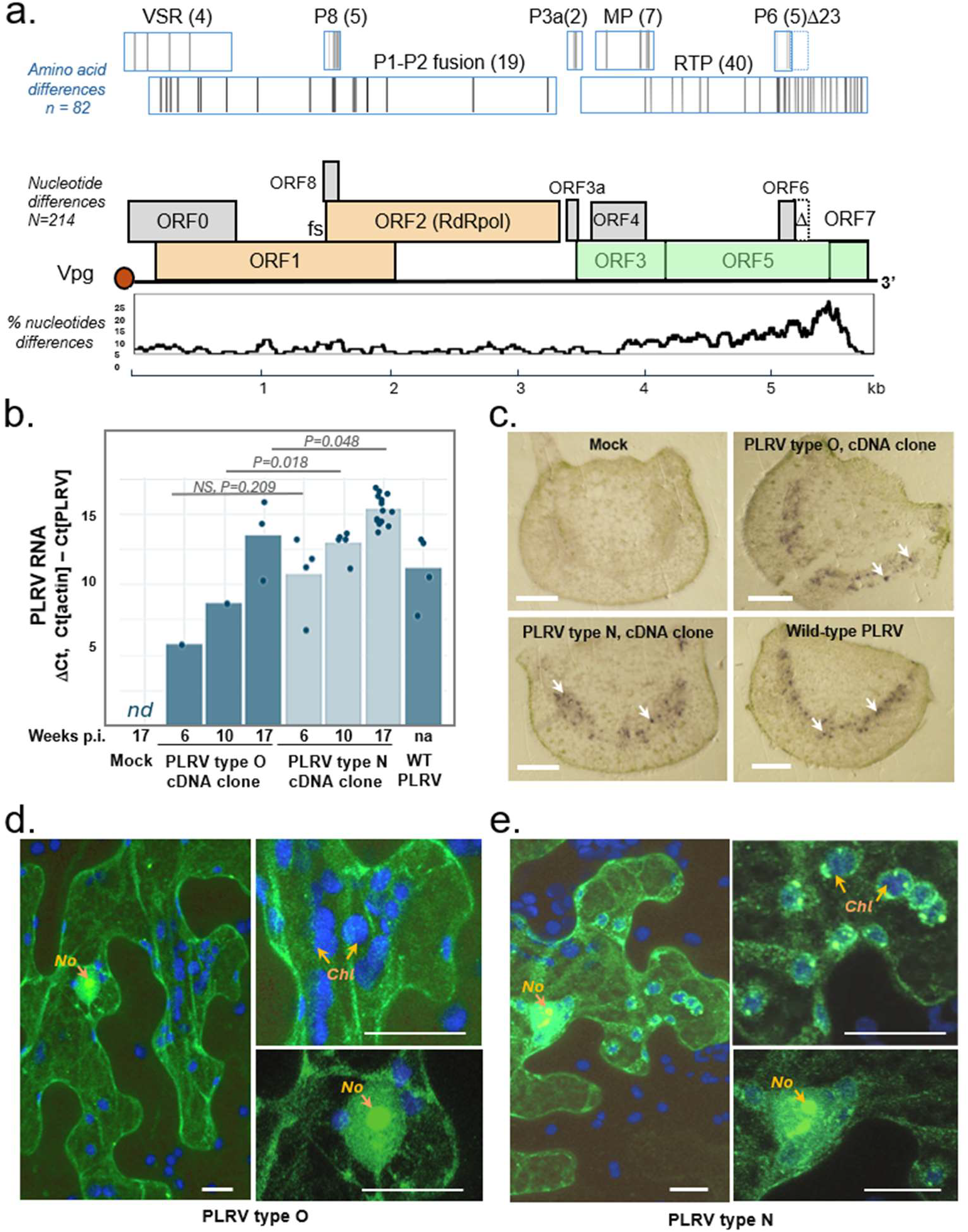
Comparative analysis of the novel and historic Scottish PLRV variants. **(a)** Organization of the PLRV genomic RNA and genetic divergence between infectious cDNA clones of the O and N types (GenBank accessions numbers OK245432 and PQ868077, respectively). The graph below PLRV genome map shows the percentage of nucleotide differences in a 100 nt sliding window. The differences between deduced AA sequences of the PLRV proteins are indicated as bars in the rectangles above the corresponding ORFs; numbers in parentheses show number of differences in a given protein; the deletion of the C-terminal 23 AAs in the P6 in the novel isolate is indicates with D. **(b)** Infection dynamics of the cDNA clone-derived PLRV O and N types in potato plants (cv Desiree),plant agroinoculated with the empty binary vector was used a negative control, the agroinfiltrated plants were sampled 6, 10 and 17 weeks post infection (wpi). The levels of PLRV genomic RNA and potato actin mRNA transcript were quantified by RT-qPCR, DCt values for the PLRV RNA normalized to the beta actin mRNA. For the plants agroinfected with the PLRV clones, results are shown only for the plants which developed PLRV infection. A potato plant, cv 18WC, naturally infected with wild-type PLRV of N type was used as a positive control. The statistical significance of the differences between PLRV levels in the plants agroinfected with the O and N type cDNA clones at the same timepoints is shown above the bars. Dots indicate individual DCt values, for 17 wpi, three samples were collected per experimental plant. **(c)** Immunoprints of potato petiole cross sections probed with polyclonal antibodies to PLRV. White arrows indicate typical PLRV infected phloem elements. Bars, 1 mm. **(d, e)** Intracellular localization of the GFP-tagged structural proteins of PLRV types O and N analyzed by confocal laser scanning microscopy. The GFP twas fused to the N-termini of the ORF3 proteins; pseudocolors: green – GFP fluorescence, blue – chloroplast autofluorescence; No – nucleolus, Chl – chloroplast; bars, 10 mm.

### 3.2. Infection dynamics of cDNA clone-derived PLRV of pylogroups O and N

We constructed a full-length infectious cDNA clone of PLRV type N using the PLRV RNA extracted from the symptomatic potato plants sourced in Scotland in 2024. The full-length genomic cDNA was cloned into the same binary vector back-bone as the one used for the construction of the PLRV clone of the Scottish phylogroup O isolate in 2021 [22]. Replication dynamics of the cloned PLRV variants was assessed in the PLRV-susceptible potato plants, cv Desiree. To inoculate potato plants with the cDNA clone-derived PLRV, fresh potato stem cuttings were incubated in the agrobacterium cultures transformed with the viral cDNA construct and subsequently rooted to produce plants as described in [22]. The PLRV levels were quantified 6, 10 and 17 weeks after inoculation by RT-qPCR using the primers designed to detect both the historic and novel PLRV types (Fig. 2b). Although agrobacterium suspensions for both PLRV cDNA clones used in agroinoculation had the same optical density, we observed a higher proportion of the plants which developed the virus infection for the N type PLRV construct (5 infected, 7 uninfected) compared with the O type PLRV clone (1 infected, 11 uninfected). The virus levels were significantly higher in the plants agroinfected with the construct of the N type compared to the O type (Fig. 3b) at 10- and 17-weeks post inoculation. Accumulation levels of both cDNA clone-derived PLRV variants in these plants (cv Desiree) were similar or exceeding those observed in the potato cv 18WC infected with a will-type PLRV (GenBank Accession PP584529).

**Figure 3.**
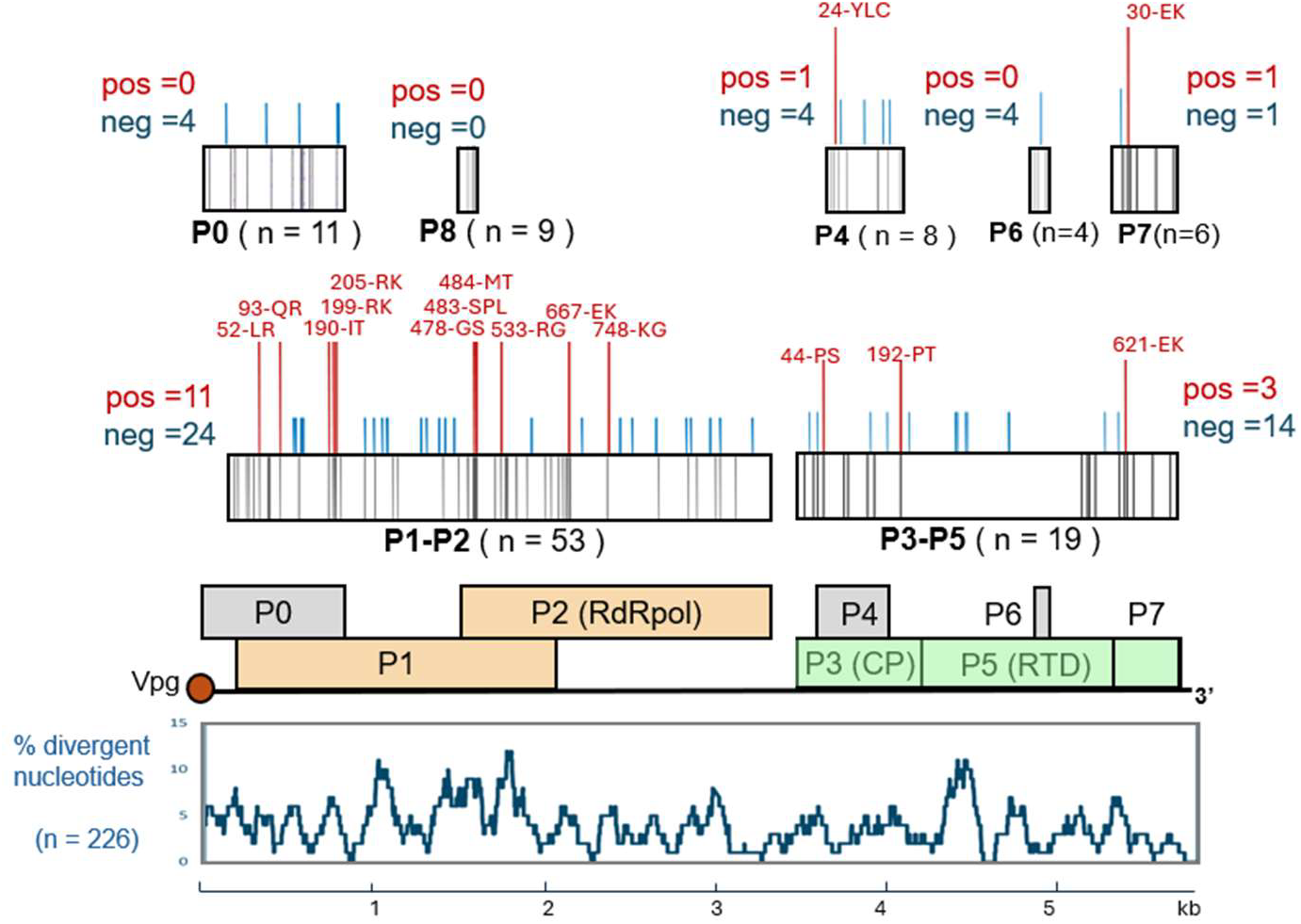
Positively and negatively selected sites in the proteins of the PLRV isolates from Scottish seed potatoes samples in 2023. Positively and negatively selected sites with posterior probability > 0.9 identified **by** FUBAR are shown as red or blue bars respectively above the boxes representing protein sequences, with amino acid changes and positions. Black bars within boxes representing protein sequences show divergent amino acid positions across all analyzed proteins (the total number of divergent amino acids for a protein is shown in brackets). The percentage of divergent nucleotide (NT) positions for a 100 NT window is shown below the map PLRV genomic RNA.

Immunoprints of the petiole cross-sections probed with antisera to PLRV were used to assess the localization of the clone-derived and wildtype PLRV in potato plants agroinoculated 17 weeks previously. Both PLRV cDNA clone-derived types, as well as wild-type PLRV were found in the phloem elements (Fig. 2c), there were no visible differences in the number and distribution of the virus infected phloem elements in the case of both cDNA clone derived and wild type PLRV (Fig. 2c), while no PLRV was detected in the control, mock inoculated potato plants.

### 3.3. Intracellular localisation of the structural proteins the historic and novel Scottish PLRV

The highest divergence between the proteins encoded by the O and PLRV types was observed for the RTP (40 of 82 divergent AAs), with most of them in the C-terminal part of the ORF5 encoded section. Information on the intracellular localization of proteins could provide clues on the protein functions. We transiently expressed the CP and RTD (encoded by the ORF3 and ORF5) block of both PLRV types with the green fluorescent protein (GFP) fused at the N-terminus of the CP in *N. benthamiana* leaves using agrobacterium-mediated infiltration approach. The plasmid constructs for the expression of the GFP-tagged PLRV proteins contained the natural amber stop codon at the end of the PLRV ORF3, which is expected to result in expression of the GFP-tagged CP (encoded by the ORF3) and the RTP (encoded by the ORF3 and ORF5). Confocal laser scanning microscopy of the epidermal and mesophyll cells of the agroinfiltrated *N. benthamiana* leaves showed that the GFP-tagged proteins were localized to the nucleus and nuclei targeting nucleoli in the case of both historic and novel PLRV variants (Fig. 2d,e), similar to previously described localization of the GFP tagged ORF3 protein of the Canadian PLRV isolate [26]. The major difference in intracellular localization between the novel and historic PLRV types was observed in the cytoplasm, where small GFP-tagged aggregates were associated with chloroplasts in the case of the novel PLRV type (Fig. 2e), while there was no chloroplast association for the phylogroup O PLRV protein (Fig. 2d). Differences in localization may indicate distinct biological properties of the structural proteins encoded by PLRV phylogroups N and O. Attachment to the chloroplast membrane was reported for many proteins of plant viruses [27,28]; however, the functional relevance of such interactions between chloroplasts and PLRV structural proteins remains to be elucidated.

### 3.4. Analysis of selection pressures acting on the PLRV genome

Phylogenetic analysis of the PLRV genomes from Scottish potato plants sampled in 2023 showed monophyletic origin and recent diversification after a bottleneck event. In total, across all 21 full-length contigs there were 3.8% of divergent NT positions (n=226), and 91 divergent AA positions in the putative P0, P1-P2, P3-P5 (RTP), P4, and P6 proteins (Fig. 3). Using FUBAR tool [20], we called 51 AA sites subjected to negative selection and 16 sites subjected to positive selection. The results of this analysis are summarized in Figure 2, which shows such sites with posterior probability exceeding 0.9. While the sites subjected to negative, stabilizing selection were distributed evenly across genome, the highest number of positively selected sites (n=8) was found in the ORF1 encoded part of the P1-P2 fusion protein. These were no positively selected sites in the ORF0 encoded protein which act as a suppressor of RNAi and interacts with the host proteins. We found three positively selected sites in the RTD protein encoded by ORF3 and ORF5. At one of these, AA position 621 in the P3-P5 fusion proteins, which correspond to the position 30 in the P7 protein, there was either lysine (K) or glutamate (E). We observed glutamate (E) at this position in higher proportion of the early field generation samples, FGs 2,3 (E/K – 6/1), compared the late filed generations, FGs 4,5,6 (E/K – 10/4), Fisher exact probability test, 2-tailed, p = 0.0237. The C-terminal part of the PLRV RTD could be involved in interaction with components of phloem and aphids, therefore aphid transmission and phloem movement may favor E621 or K621 respectively.

### 3.5. Possible impact of transmission routes on PLRV diversification

Analysis of *de novo* contigs produced for the libraries showed variation in the divergency levels in the PLRV genome regions containing the structural and non-structural genes within FGs (Fig. 4a). We observed that in early FGs the ratio between the numbers of divergent NTs in the non-structural genes section and in the structural genes section (20 to 1 in FG2, 68 to 16 for FG3), was higher than in the late FGs (88 to 36 for FG5, 89 to 43 for FG6), Fig. 4a). Contingency table analysis showed that these differences were statistically significant between early and late FGs. For FG6, the proportion of the divergent positions in the structural genes section was significantly higher compared to those in FG2 and FG3 (p = 0.008 and p = 0.040, correspondingly). Significant difference in relative distribution of divergent NTs was also observed between FG2 and FG5 (p = 0.027), Fig. 4b.

**Figure 4.**
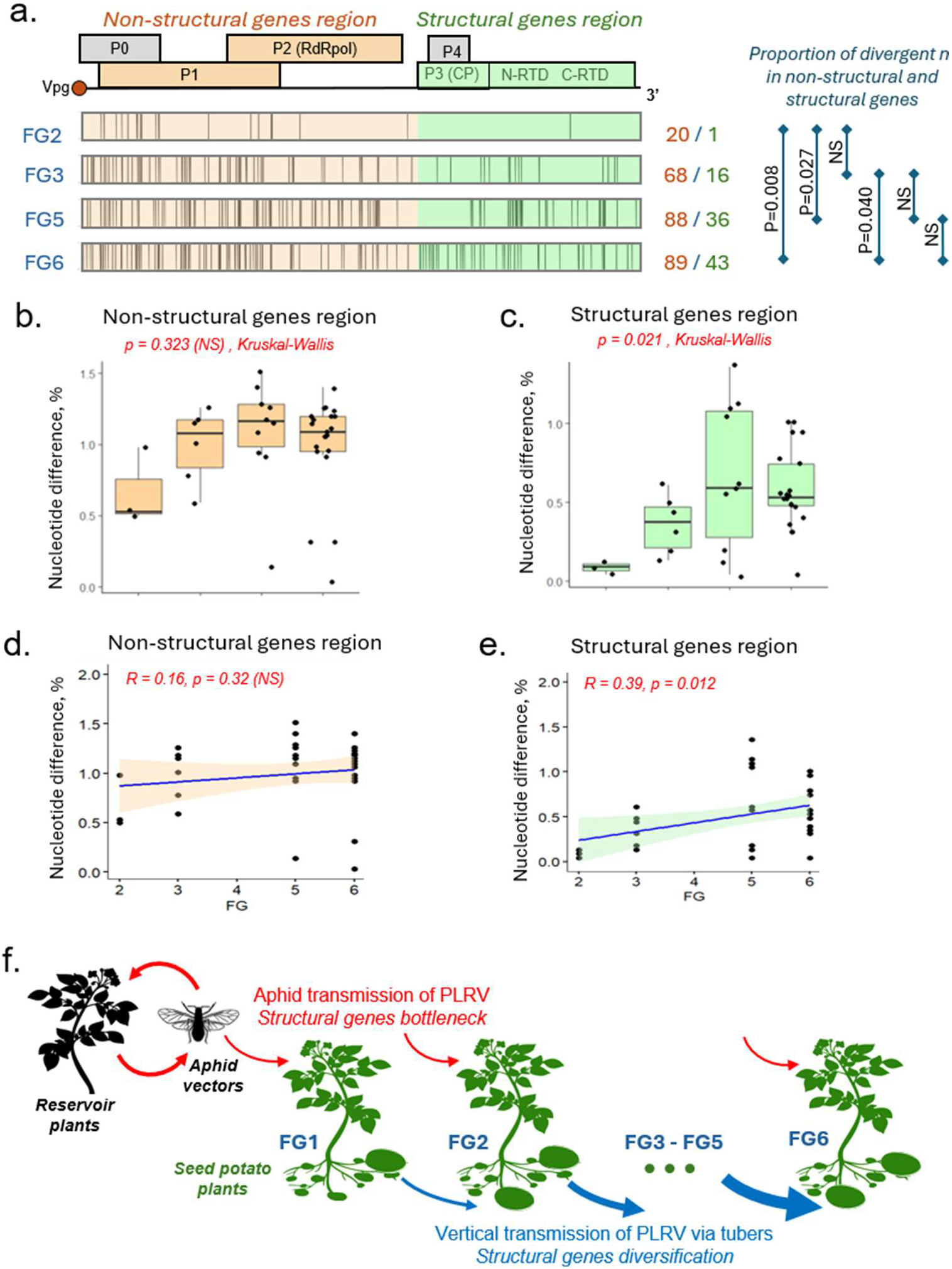
Analysis of selection pressures on PLRV genome imposed by the aphid and tuber transmission routes. **(a)** Distribution of divergent nucleotide (NT) positions, represented as vertical bars, in the PLRV genomes within the same FG for the de novo contigs. The schematic representation of PLRV genome is shown above. The numbers of divergent NTs in the genome sections for FGs (non-structural section /structural section), and the significance of the differences of the divergent NT proportions in the non-structural and structural sections assessed by contingency table analysis are shown at the right side. **(b-e)** Analysis of PLRV divergence within FGs for the de novo contigs. Boxplot shows pair-wise NT divergence, calculated for the sections of the PLRV genome: **(b)** non-structural genes region (1 - 3.5 kb region), **(c)** structural genes region (3.5 - 5.9 kb region). Dots represent the values for individual pairwise comparisons between libraries within each of FGs; **(d**,**e)** Scatter plots for the pairwise NT divergence values within FG. Shown are Pearson correlation coefficient (R) and significance level. **(f)** Proposed model of PLRV circulation. Plants of the first field generation (FG1) are grown from virus-free tubers, the progeny tubers are planted next season to produce FG2. The cycle continues up to FG6. Early FGs are infected by aphids, later FGs could be to be infected with PLRV received through tuber from infected parental plants. Red arrows indicate aphid transmission, blue arrows – vertical transmission via tubers.

We calculated pairwise percentages of nucleotide divergence between de novo contigs within n FGs 2, 3, 5 and 6, separately for the PLRV genome sections containing non-structural genes (positions 1 – 3.5 kb), and (Fig. 4cd). We observed significant differences of the structural gene section in the late FGs (Kruskal-Wallis test, c^2^ = 9.7271, df = 3, p-value = 0.0210), while the percentages of nucleotide divergence for the non-structural gene section (5’ proximal section part of the genomic RNA, 1-35 kb) were not significantly different across all FGs (Kruskal-Wallis test, c^2^ = 4.3999, df = 3, p = 0.2214). Our analysis showed a significant increase of diversity of the structural gene block in the later FGs (p < 0.05), while diversity of the non-structural genes did not show a significant change across FGs (Fig. 4e,f).

Based on these results we suggest a hypothetical model of the PLRV transmission in seed potatoes, and the impact of transmission routes on genetic diversity of PLRV (Fig. 4f). Potato plants could be infected with PLRV by aphids or could acquire PLRV via tuber transmission from infected mother plants. FG1 plants are infected only by aphids, but plants of later FGs could be infected either by aphid vectors (horizontal transmission), or through tuber propagation (vertical transmission). Reduced diversity of the structural genes in the early FG2 and FG3, which are more likely to receive the virus from aphids rather than from tubers compared to the later FG5 and FG6, may suggest that the aphid transmission imposes a genetic bottleneck acting on the genes of structural proteins involved in interaction with aphids.

## 4. Discussion

Epidemics of vector-borne plant viruses could have a significant economic impact by affecting yield and quality of crops, but their causes are not fully understood [14]. Several factors could contribute to the increase of virus incidence, of which introduction and emergence of novel virus variants, and changes in insect vector populations are the most important drivers [29].

In this study we analyzed genetic diversity of PLRV circulating in Scottish potato crops. High-throughput sequencing of PLRV genomes from potato samples sourced in the years 2023 and 2024 showed that the N phylogroup of PLRV has displaced phylogroup PLRV isolates of phylogroup O, which was prevalent in Scottish potatoes since at least 1989 (Fig. 1) [2, 22]. All Scottish PLRV isolates sourced in 2023-2024 showed highest similarity with the PLRV isolate identified in Germany in 2012, which was reported as causing asymptomatic infection in some potato cultivars in glasshouse conditions [24]. Although all Scottish 2023-2024 PLRV isolates of the N phylogroup sequenced in this study were identified in field-grown symptomatic potato plants, there is a possibility that different pathogenic characteristics of the N type and/or other environmental or cultural factors might have contributed to selection and prevalence of the N phylogroup [30].

Sequence analysis showed that the RNA genomes of Scottish PLRV isolates of the O and N phylogroups have approximately 4% of divergent NTs (Fig. 2a). Comparison of the proteins encoded by the infectious cDNA clones of these types (Fig. 1, pointed by arrows) showed 82 divergent AA positions in all proteins, with the largest number located in the C-terminal part of the RTP and the P7 (Fig. 2a). Another feature of the isolates belonging to the PLRV type N phylogroup is a truncation of P6 due to NT substitution leading to introduction of a stop codon resulting in the deletion of 23 C-terminal AAs of the originally 62 AA-long protein.

These AA difference may lead to the alteration of biological properties of the virus isolated. For example, the RTP is a minor component of the PLRV capsid which interacts with both plant and aphid components [5,9]. Therefore, changes in RTP may affect virus movement within plants and possibly the aphid transmission. Indeed, comparison of infection dynamics of the cDNA clone derived PLRV in potato plants suggested that the PLRV clone of the N type replicated to a higher level compared to the O type (Fig. 2b). Further studies in natural conditions of transmission and plant growth are needed to confirm these preliminary observations. Also, localisation studies showed that structural proteins of the N and O PLRV types have some differences in intracellular localisation (Fig. 2d,e). The P0, P1 and P7 proteins of PLRV involved in changing attractiveness of aphids to infected plants by affecting aphids’ fecundity via modulation of phytohormone production in *N. benthamiana* plants [25], therefore differences between the novel and historic PLRV type may have an effect on the virus ability to be transmitted by the aphid vectors.

Phylogenetic analysis of PLRV from infected seed potatoes sampled in 2023 showed monophyletic origin (Fig. 1) making it possible to analyze selective pressures action on the PLRV genomes. In total, 51 AA sites subjected to negative selection and 16 sites subjected to positive selection in the P0, P1-P2 fusion, P3-P5 (RTP), P4, and P6 proteins (Fig. 3). We found no positively selected sites in the P0 protein, which act as a suppressor of antiviral RNAi. The action of the P0 involves interaction with the host S-phase kinase-related protein-1 which is required to direct ubiquitination and degradation of Argonaute1 [31]. Such interaction requires adaptation to a specific plant host, therefore the lack of diversification of the P0 may indicate that circulation of PLRV involves solely potato plants. Positively selected sites in the P3 and RTP, as well as in the P7, such as position 621 in the RTP (Fig. 3), may reflect adaptation of PLRV to the phloem transmission following aphid transmission step.

PLRV is transmitted from virus-infected to the virus-free plants by aphids, principally by *Myzus persicae*, which vector the virus in non-replicative circulative manner [4,5]. Tuber propagation of potatoes also allows transmission of PLRV from infected plants to the next vegetative generations [32]. Seed potato production starts with planting virus-free mini tubers; from which the progeny tubers are subsequently propagated over several FGs. Therefore, all PLRV infected plants of FG1 are infected only via aphid transmission, while the later FGs could be infected either by the aphids or could have acquired PLRV vertically form parental infected plants via infected tubers (Fig. 4f). In this study we investigated the impacts of vertical (tuber) and horizontal (aphid) transmission on genetic diversity of PLRV via high-throughput sequencing of PLRV from different FGs of seed potatoes. Plants of early FGs (FG2 and FG3) are more likely to be directly infected with PLRV by the aphid vectors; while the late FGs (FG4 to FG6) were likely to be infected with the virus propagated via tuber transmission which underwent up to five field generations.

We found that recently aphid-transmitted PLRV variants from FG2 and FG3 had significantly lower genetic diversity of 3’ genome section coding for the structural proteins compared to those from the late field generations, while diversity levels of the non-structural gene block were not significantly different across all FGs, from FG2 to FG6 (Fig. 4a-e). This suggests that aphid transmission may impose a strong genetic bottleneck on the structural genes of PLRV, leading to the reduction of diversity. This could be driven by selection of the PLRV variants with the structural genes better adapted to aphid transmission. The structural genes showed diversification in the later FGs, which might be a result of the absence of the need to be adapted to the aphid transmission and/or possible adaptation to the tuber transmission (Fig. 4a-e). Notably, the positively selected divergent aa 621 in the RTP which showed significant association with filed generation (glutamate in FG2, 3, and lysine in FG4,5,6) is located in the region of RTD, mutation in which affects aphid transmission, possibly by affecting passage through the aphid gut membrane [33,34]. We hypothesized that transmission of PLRV via tubers for several vegetative generations (FGs) in the absence of the aphid transmission events may result in accumulation of the genetic variants of PLRV with decreased adaptation to aphid vectoring and/or higher adaptation to vertical transmission (Fig. 4f).

The lack of genetic bottleneck associated with the aphid transmission which acted on the non-structural genes (such as ORF0, 1 and 2) was in the agreement with the non-replicative nature of the aphid transmission of the virus and the lack of interaction of the replicative genes of PLRV with the aphid components. The impact of such diversification of PLRV on the virus pathogenicity and ability to be transmitted by aphid vectors, and possible implications for the virus epidemiology require further investigation.

## 5. Conclusions

We identified a prevalent PLRV phylogroup in Scottish potatoes crops which, by 2023-2024 had displaced a different phylogroup present for over four decades. The currently prevalent PLRV type is phylogenetically related to a German isolate identified in 2012. Analysis of PLRV populations from different field generations of seed potato showed reduction of genetic diversity of the structural genes in PLRV recently transmitted by aphids, which suggest a strong genetic bottleneck acting on the structural genes of PLRV.

## Supplementary Materials

**Supplementary Figure S1.**
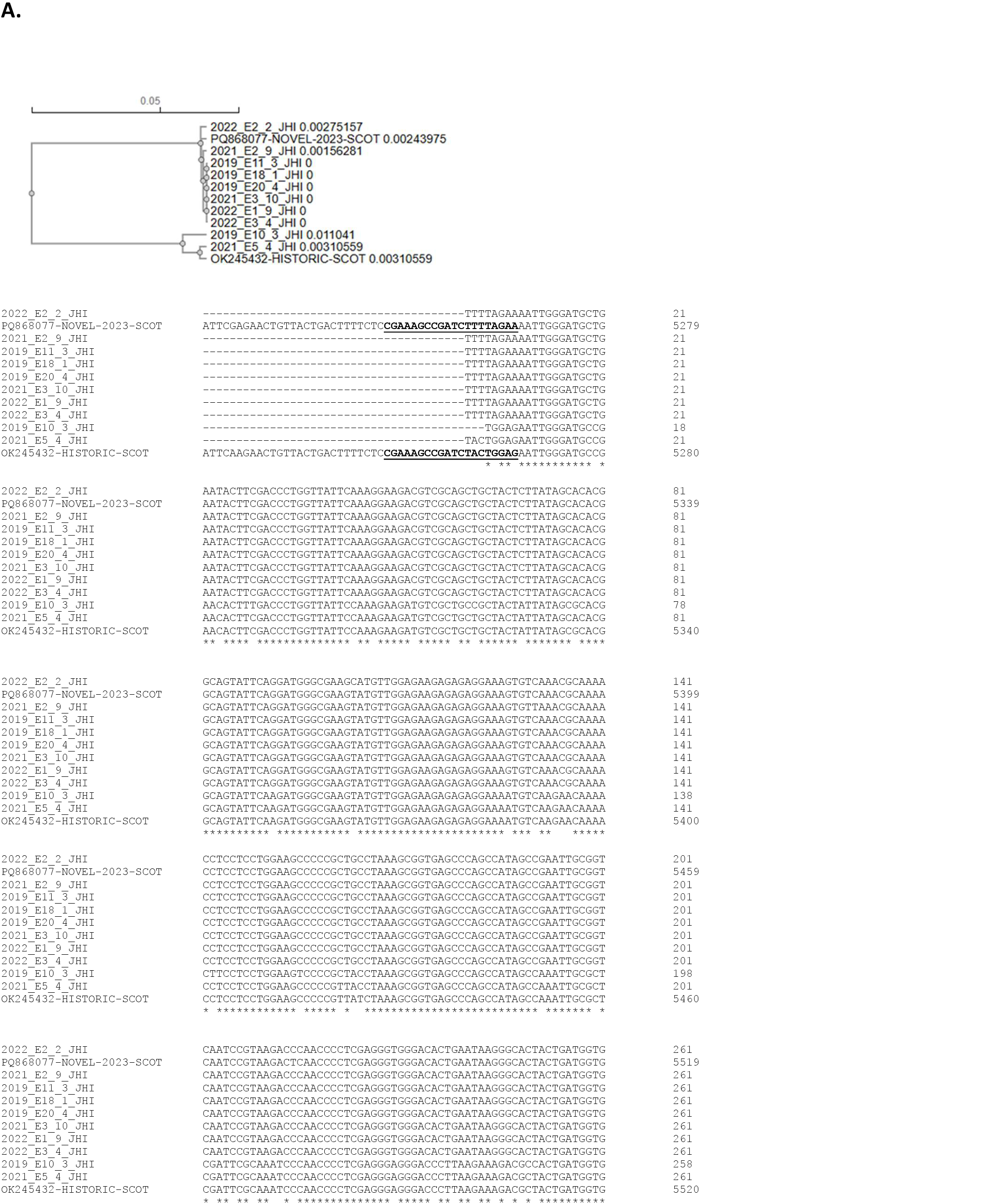
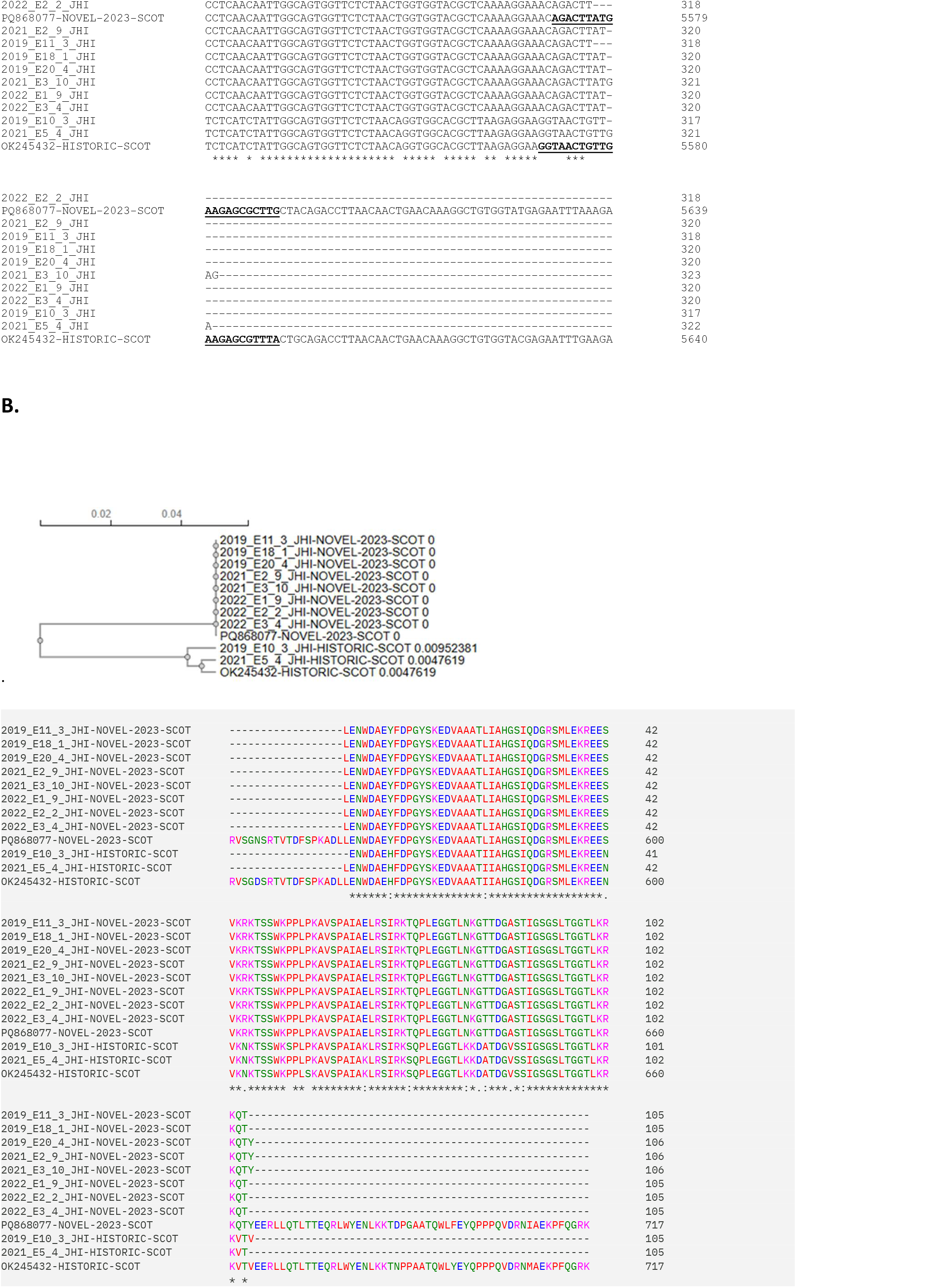
Identification of PLRV types O and N in Scottish potato samples collected in 2019 – 2022. **A**. Phylogeny and alignment of the partial nucleotide sequences of the 3’ sections of the Scottish isolates, year of collection indicated by prefix in the sequence ID; also included the corresponding regions of the PLRV type O (GenBank Accession OK245432), and the PLRV type N (GenBank Accession PQ868077). **B**. Phylogeny and alignment of the partial ORF5 protein sequences encoded by the 2019-2022 Scottish PLRV isolates.

## Author Contributions

Conceptualization, EVR; methodology, EVR, GHC., CL, CT; validation, formal analysis, EVR, GHC, CL; investigation, GHC, CT, CL, EVR; resources, CL, LT; data curation, EVR, CL; writing - original draft preparation, EVR; writing- review and editing, EVR, GHC, LT, CL; visualization, EVR; supervision, EVR, CL; project administration, EVR, CL; funding acquisition, EVR, LT, CL. All authors have read and agreed to the published version of the manuscript.

## Data Availability Statement

The nucleotide sequence of PLRV isolates produced in this study are deposited to GenBank, accession numbers PP584529 - PP584549, PQ047449 - PQ047454, PV647837, PV647838.

## Acknowledgments

We thank Eric Anderson and Zach Reilly (Scottish Agronomy Ltd.), Archie Gibson (Agrico UK Ltd.) for providing potato samples, and Pete Hedley for high throughput sequencing, and Philip Greenspoon for his helpful discussions and suggestions on virus population comparison.

## References

1. Harrison, B.D. Potato leafroll virus. Descriptions of plant viruses 1984, 291. Kew CMI/AAB

2. Mayo, M.A.; Robinson, D.J.; Jolly, C.A.; Hyman, L. Nucleotide Sequence of Potato Leafroll Luteovirus RNA. J. Gen. Virol. 1989, 70, 1037–1051. 10.1099/0022-1317-70-5-1037

3. Somera, M.; Fargette, D.; Hébrard, E.; Sarmiento, C. ICTV Report Consortium. ICTV virus taxonomy profile: Solemoviridae. J. Gen. Virol. 2021, 102, 001707. 10.1099/jgv.0.001707

4. Taliansky, M.; Mayo, M.A.; Barker, H. Potato leafroll virus: A classic pathogen shows some new tricks. Mol. Plant Pathol. 2003, 4, 81–90. 10.1046/j.1364-3703.2003.00153.x

5. Farooq, T.; Hussain, M.D.; Shakeel, M.T.; Riaz, H.; Waheed, U.; Siddique, M.; Shahzadi, I.; Aslam, M.N.; Tang, Y.; She, X.; He, Z. Global genetic diversity and evolutionary patterns among Potato leafroll virus populations. Front. Microbioll. 2022, 13:1022016. https://www.frontiersin.org/journals/microbiology/articles/10.3389/fmicb.2022.1022016/full

6. Killick, R.J.; The effect of infection with potato leafroll virus (PLRV) on yield and some of its components in a variety of potato (Solanum tuberosum). Ann. Appl. Biol. 1979, 91, 67–74

7. Marshall, B.; Barker, H.; Verral, S.R. Effects of potato leafroll virus on the crop processes leading to tuber yield in potato cultivars which differ in tolerance of infection. Ann. Appl. Biol. 1988, 113, 297–305.

8. DeBlasio, S.L.; Johnson, R.; Sweeney, M.M.; Karasev, A.; Gray, S.M.; MacCoss, M.J.; Cilia, M. The Potato leafroll virus structural proteins manipulate overlapping, yet distinct protein interaction networks during infection. Proteomics 2015, 15, 2098–2112. http://dx.doi.org/10.1002/pmic.201400594

9. Schiltz, C.J.; Wilson, J.R.: Hosford, C.J.; Adams, M.C.; Preising, S.E.: DeBlasio, S.L., MacLeod, H.J.; Van Eck, J.; Heck, M.L.; Chappie, J.S. Polerovirus N-terminal readthrough domain structures reveal molecular strategies for mitigating virus transmission by aphids. Nature Communications 2022. 13, 6368 10.1038/s41467-022-33979-2

10. Peter, K.A.; Liang, D.; Palukaitis, P.; Gray, S.M. Small deletions in the potato leafroll virus readthrough protein affect particle morphology, aphid transmission, virus movement and accumulation. J. Gen. Virol. 2008, 89, 2037–2045. 10.1099/vir.0.83625-0

11. Olmedo-Velarde, A.; Wilson, J.R.; Stallone, M.; DeBlasio, S.L.; Chappie, J.S.; Heck, M. Potato leafroll virus molecular interactions with plants and aphids: Gaining a new tactical advantage on an old foe. Physiological and Molecular Plant Pathology 2023, 125, 102015. 10.1016/j.pmpp.2023.102015

12. Skelsey, P. Landscape-scale patterns and predictors of potato viruses in Scotland. Plant Pathology, 2024, 73, 1553–1572. 10.1111/ppa.13891

13. Whatty, M.; Collier, R. The Impact of the EU Neonicotinoid Ban on UK Carrot and Lettuce Production and Prospects for Future Aphid Management. Outlooks on Pest Management, 2023, 34, 20–29. 10.1564/v34_feb_07

14. Jones, R.A.C. Global Plant Virus Disease Pandemics and Epidemics. Plants 2021, 10, 233. 10.3390/plants10020233

15. Davie, K.; Holmes, R.; Pickup, J.; Lacomme, C. Dynamics of PVY strains in field grown potato: Impact of strain competition and ability to overcome host resistance mechanisms. Virus Res. 2017, 241, 95–104 10.1016/j.virusres.2017.06.012

16. Antipov, D.; Raiko, M.; Lapidus, A.; Pevzner, P.A. METAVIRALSPADES: assembly of viruses from metagenomic data. Bioinformatics 2020, 36, 4126–4129. 10.1093/bioinformatics/btaa490

17. Altschul, S.F.; Gish,W.; Miller,W.; Myers, E.W.; Lipman, D.J. Basic local alignment search tool. J. Mol. Biol. 1990, 215, 403–410. 10.1016/S0022-2836(05)80360-2

18. Sievers, F.; Higgins S.G. Clustal omega. Current protocols in Bioinformatics 2014, 10.1002/0471250953.bi0313s48

19. Felsenstein, J. PHYLIP - Phylogeny Inference Package (Version 3.2). Cladistics 1989, 5, 164–166. http://evolution.genetics.washington.edu/phylip.html

20. Weaver, S.; Shank, S.D.; Spielman, S.J.; Li, M.; Muse, S.V.; Kosakovsky Pond, S.L. Datamonkey 2.0: A Modern Web Application for Characterizing Selective and Other Evolutionary Processes. Mol. Biol. Evol. 2018, 35, 773–777. 10.1093/molbev/msx335

21. R Development Core Team. R: A language and environment for statistical computing. R Foundation for Statistical Computing, Vienna, Austria. 2009, ISBN 3-900051-07-0, URL http://www.R-project.org.

22. Cowan, G.; MacFarlane, S.; Torrance. L. A new simple and effective method for PLRV infection to screen for virus resistance in potato. J. Virol. Meth. 2023, 315, 114691. 10.1016/j.jviromet.2023.114691

23. Xiang, C.; Han, P.; Lutziger, I.; Wang. K; Oliver D.J. A mini binary vector series for plant transformation. Plant Molecular Biology 1999, 40, 711–717. https://link.springer.com/article/10.1023/a:1006201910593

24. Hühnlein, A., Schubert, J., Zahn, V., & Thieme, T. 2016, Examination of an isolate of potato leaf roll virus that does not induce visible symptoms in the greenhouse. European Journal of Plant Pathology, 145(4), 829–845. 10.1007/s10658-016-0872-3.

25. Patton, M.F.; Bak, A.; Sayre, J.M.; Heck, M.L.; Casteel, C.L. A polerovirus, Potato leafroll virus, alters plant-vector interactions using three viral proteins. Plant Cell Environ. 2020, 43, 387–399. https://onlinelibrary.wiley.com/doi/full/10.1111/pce.13684

26. Haupt, S.; Stroganova, T.; Ryabov, E.; Kim, S.H.; Fraser, G.; Duncan, G.; Mayo, M.A.; Barker, H.; Taliansky, M. Nucleolar localization of potato leafroll virus capsid proteins. J. Gen. Virol. 2005, 86, 2891–2896. https://www.microbiologyresearch.org/content/journal/jgv/10.1099/vir.0.81101-0

27. Bhattacharyya, D.; Chakraborty, S. Chloroplast: The Trojan horse in plant–virus interaction. Mol. Plant Pathol. 2018, 19, 504–518. https://bsppjournals.onlinelibrary.wiley.com/doi/10.1111/mpp.12533

28. Zhao, J.; Zhang, X.; Hong, Y.; Liu, Y. Chloroplast in plant-virus interaction. Front. Microbiol. 2016, 7, 1565. https://www.frontiersin.org/journals/microbiology/articles/10.3389/fmicb.2016.01565/full

29. Anderson, P.K.; Cunningham, A.A.; Patel, N.G.; Morales, F.J.; Epstein, P.R.; Daszak, P. Emerging infectious diseases of plants: Pathogen pollution, climate change and agrotechnology drivers. Trends Ecol. Evol. 2004, 19, 535–544. https://www.cell.com/trends/ecology-evolution/fulltext/S0169-5347(04)00218-6

30. Kumar, R.; Tiwari, R.K.; Sundaresha, S.; Kaundal, P.; Raigond, B. Potato Viruses and Their Management. In Sustainable Management of Potato Pests and Diseases; Springer: Singapore, 2022; pp. 309–335.

31. Pazhouhandeh, M.; Dieterle, M.; Marrocco, K.; Lechner, E.; Berry, B.; Brault, V.; Hemmer, O.; Kretsch, T.; Richards, K.E.; Genschik, P.; et al. F-box-like domain in the Polerovirus protein P0 is required for silencing suppressor function. Proc. Natl. Acad. Sci. USA 2006, 103, 1994–1999. 10.1073/pnas.0510784103

32. Sastry, K.S. (2013). Plant Virus Transmission Through Vegetative Propagules (Asexual Reproduction). In: Seed-borne plant virus diseases. Springer, India. 10.1007/978-81-322-0813-6_9

33. Jolly, C.; Mayo, M. Changes in the amino acid sequence of the coat protein readthrough domain of potato leafroll luteovirus affect the formation of an epitope and aphid transmission. Virology 1994, 201, 182–185. 10.1006/viro.1994.1283

34. Rouze-Jouan, J.; Terradot, L.; Pasquer, F.; Tanguy, S.; Ducray-Bourdin, D.G. The passage of Potato leafroll virus through Myzus persicae gut membrane regulates transmission efficiency. J. Gen. Virol. 2001, 82, 17–23. 10.1099/0022-1317-82-1-17

